# Idiosyncrasy in gestural communication: a case study of hand-clapping in a Barbary macaque (*Macaca sylvanus*)

**DOI:** 10.1101/2024.09.09.611981

**Authors:** Tiffany Claire Bosshard, Marie Hirel, Hélène Meunier, Julia Fischer

**Author notes:** Corresponding author: Tiffany Claire Bosshard, Address: German Primate Center (DPZ), Kellnerweg 4, 37077 Göttingen, Germany. The study was purely observational. It was done on monkeys housed in a tourist park; therefore, the intrusion by the observers can be considered minimal. Video supporting our observations are part of this work.

## Abstract

While it is well established that apes invent or individually learn new gestures, cases of development and use of novel gestures in monkeys are more rarely described. We report a case of a novel, idiosyncratic gesture in a Barbary macaque (*Macaca sylvanus*) at ‘La Forêt des Singes’, Rocamadour, France. One adult male, Jomanix, was observed hand-clapping. To our knowledge, hand-clapping has never been described before in this species. To hand-clap, the male briefly shifted his weight onto his legs, lifted his upper body, and clapped both hands together. We recorded 30 instances of hand-clapping. Twenty-five of these hand-claps occurred in combination with other agonistic signals, such as lunges and open mouth threats. Recipients either responded with counter-aggression (*N* = 9) or a submissive response (*N* = 16). In five of the 30 events, the context was unclear. These observations suggest that the gesture constitutes an agonistic signal. According to the staff at ‘La Forêt des Singes’, the hand-clapping may have been copied from staff members who occasionally hand-clap to shoo the animals away from areas where they were not supposed to be, but that notion remains speculative. In the meantime, another subject from the same group reportedly started to hand-clap, but the subject had passed away before we could document the behaviour. The observations show that Jomanix can flexibly combine a novel gesture with other established communicative signals. The hand-clap is goal-directed and fulfils the criteria for first-order intentional communication. This case, as well as anecdotal reports from a Tonkean macaque (*Macaca tonkeana*) hand-clapping to get attention, reveals greater flexibility in the gestural communication of this genus than previously assumed but also underscores that social learning of the production of communicative gestures occurs rarely in this taxon.

## 1. INTRODUCTION

Manual and bodily gestures represent a central component of the signal repertoire that nonhuman primates (hereafter primates) use to communicate (Call & Tomasello, 2007; Cartmill et al., 2012; Pika & Liebal, 2012). Gestural communication occurs within various social contexts, such as grooming (de Waal, 1988), mating (Genty & Zuberbühler, 2014), social play (McCarthy et al., 2013), food sharing (Fröhlich et al., 2017), joint travel (Fröhlich et al., 2016), or aggression (Scott, 2013). Yet, the mechanisms by which primates acquire novel gestures remain a subject of debate. There are several theories regarding the emergence and development of gestures in primates, which broadly follow the lines of gestures being either genetically determined, socially acquired, or subject to an interplay of both processes (reviewed in Liebal et al., 2019). However, one reoccurring observation across studies is the presence of idiosyncratic gestures, i.e., gestures produced only by single individuals within a group (Tomasello et al., 1994). While not always given much importance, the occurrence of idiosyncratic gestures serves as an opportunity to elucidate the outer boundaries of a species’ reaction norm.

It is well established that apes can invent or individually learn new gestures, and gestural idiosyncrasy was reported for all great ape species (e.g., chimpanzees, *Pan troglodytes*, Roberts et al., 2014; Tomasello et al., 1994; bonobos, *Pan paniscus*, Pika et al., 2005; gorillas, *Gorilla g. gorilla*, Byrne & Tanner, 2006; Genty et al., 2009; Pika et al., 2003; orang-utans, *Pongo abelii* and *pygmaeus*, Amici & Liebal, 2023; Cartmill & Byrne, 2010). Cases of development and use of novel gestures in monkeys, in contrast, are more rarely described. For instance, a hamadryas baboon (*Papio hamadryas*) was observed swinging one hand towards a partner in a waving gesture (Dube, 2013), a juvenile olive baboon (*Papio anubis*) was observed holding a part of his upper arm between his teeth and shaking the rest of the arm in a trunk-like gesture during play (Molesti et al., 2020), and a Barbary macaque (*Macaca sylvanus*) was observed using his hand to conceal his facial expressions during playful and aggressive interactions (Thunström et al., 2014). Taking into account group-specific novel gestures among monkeys, mandrills (*Mandrillus sphinx*) were reported displaying a new gesture of hand extension used in social contexts (Laidre, 2008), and bonnet macaques (*Macaca radiata*) were reported stretching out their hands and ‘begging’ humans for food (Sinha, 2005).

We here report a case of a novel idiosyncratic gesture in a Barbary macaque, namely one adult male, Jomanix, who was observed hand-clapping. To hand-clap, the male briefly shifts his weight onto his legs, lifts his upper body, and claps both hands together (Figure 1). The gesture is well-audible beyond ten meters. To our knowledge, this is a new gesture in the species’ behavioural repertoire (Hesler & Fischer, 2007), at least at this location. Over more than 30 years of research from our group at this facility, we had never observed it before.

**Figure 1.**
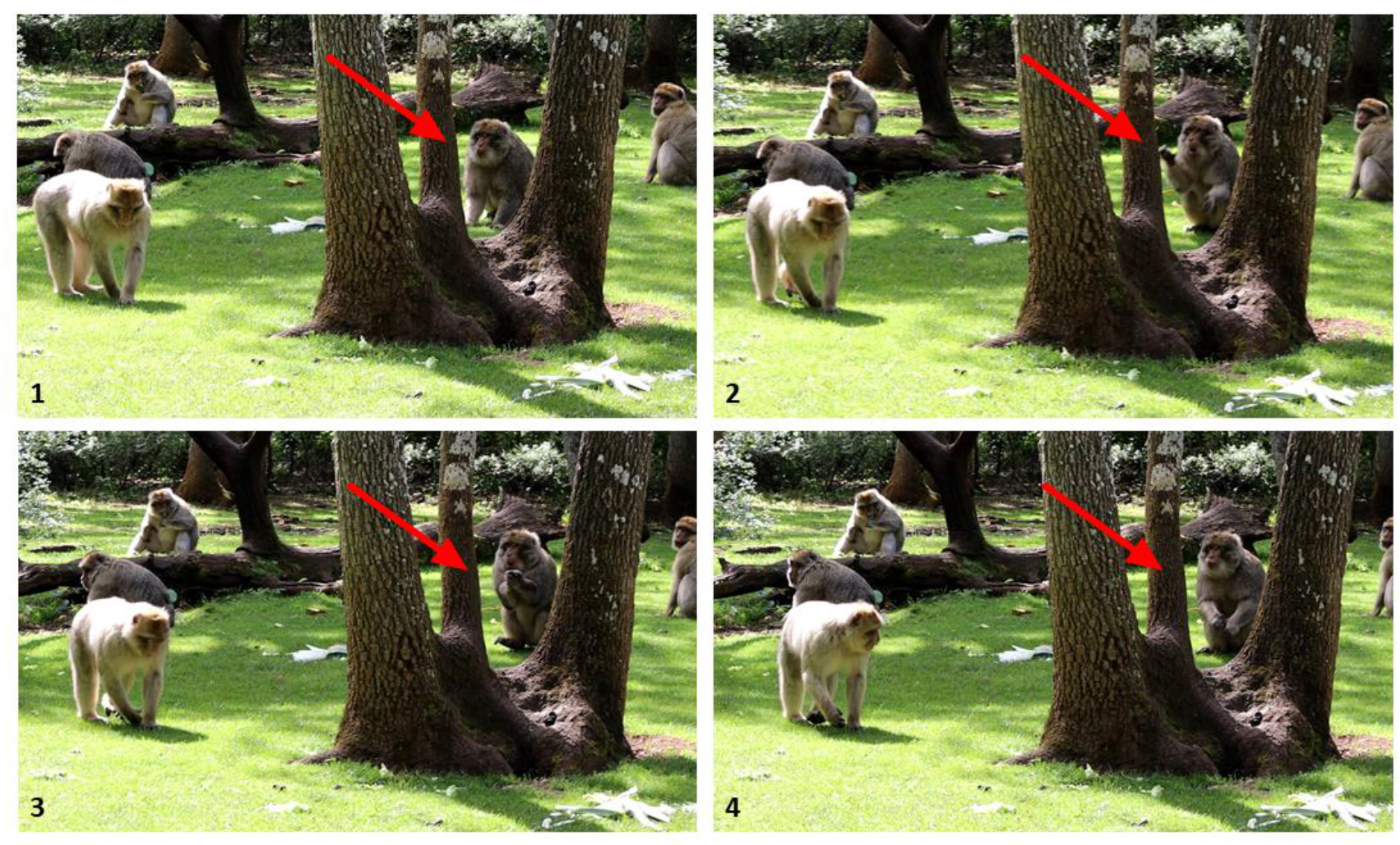
Video still sequence of Jomanix hand-clapping (red arrows), numerically ordered.

We aimed to determine whether Jomanix uses the hand-clap within the realm of gestural communication, and if so, which function the gesture may have. To attain communicative function, gestures need to occur in specific contexts, and be directed at a specific individual with the purpose of influencing their behaviour, i.e., evoke a measurable response (Fischer & Price, 2017; Smith & Harper, 2003). We additionally report another case of hand-clapping in a Tonkean macaque (*Macaca tonkeana*) male; this report is largely anecdotal but informative to evaluate the scope for social learning of gestures in this genus.

## 2. METHODS

### 2.1 Study sites and subjects

We carried out behavioural observations in a population of Barbary macaques housed at ‘La Forêt des Singes’, in Rocamadour, France, between March and June of 2022 and 2023. This park consists of a 20-hectare enclosed forest in which the animals range freely and live outdoors year-round (de Turckheim & Merz, 1984). Throughout the months of April to November, the park is open to visitors, who can watch the monkeys from designated walking paths. The monkeys are thus well-habituated to human observers, yet have the opportunity to distance themselves from people if they wish to do so. While the animals feed on the naturally available vegetation in the forest, they are also provided with fruits, vegetables, and grains several times per day to supplement their diet. Water is available *ad libitum*. At the time of the observations, there was a total of 146 monkeys housed in the park, split into three naturally-formed groups of more or less 50 individuals each.

Jomanix is part of a group which, at the time of the first observations, consisted of 46 individuals, including 27 males (25 adults, two juveniles under the age of three) and 19 females (18 adults, one juvenile). Jomanix was then 15 years old (i.e., born in 2007) and was, at least at that time, the dominant male of the group (normalised David’s scores from 2022). According to the park staff, he became the dominant male of his group in 2019.

The case of hand-clapping in the Tonkean macaque occurred at the Centre de Primatologie – Silabe de l’Université de Strasbourg, in France. The Tonkean macaques at the facility have permanent access to a wooded outdoor enclosure of 3788m^2^, connected to an indoor space of 20m^2^. The animals are fed with commercial primate pellets every day, and fresh fruits and vegetables once a week. They have access to water *ad libitum*. We were there in 2018 for a study on appearance-reality discrimination in Tonkean macaques and brown capuchins (*Sapajus apella*) (Hirel et al., 2020), and in 2023 for a study on social evaluation in the same species (Hirel et al., in prep.). The hand-clapping individual, Wallace, was 11 years old at the time of the first study (i.e., born in 2007), and part of a group consisting of 28 individuals (16 males, 12 females). In 2019, Wallace was moved to an only-male group of six individuals living in a wooded outdoor enclosure of 1364m^2^ with an indoor space of 10m^2^. The Tonkean macaques are highly habituated to the presence of human experimenters and have willingly participated in several cognitive studies.

### 2.2 Data collection

From March to June 2022, we conducted behavioural observations of all adult individuals in Jomanix’s group as part of a study on social responsiveness in Barbary macaques. During this time, we first observed the hand-clapping behaviour displayed by Jomanix, which we then included in our observations in addition to the established Barbary macaque ethogram (Hesler & Fischer, 2007). Observations consisted of thirty-minute continuous focal samplings, during which we recorded the animals’ general activity, as well as all affiliative and agonistic interactions. Agonistic interactions were also recorded *ad libitum* for the purpose of establishing the dominance hierarchy. We collected an average of 40 focal observations per individual, resulting in approximately 20 hours of observation of Jomanix. From March to June 2023, we conducted focal behavioural observations, as described above, on monkeys of a different group but visited Jomanix’s group when time permitted for opportunistic recordings of further hand-clap events. In May of 2024, we returned to the park to teach a two-week practical course and again recorded hand-clap events when possible.

In 2022, we collected behavioural data using handheld devices (GIGASET GX290 plus) equipped with Pendragon Forms (Pendragon Software Cooperation), and video-recorded hand-clap events whenever feasible. In 2023 and 2024, we recorded the hand-claps on video only. Across the three observation periods, we recorded 30 hand-clap events from Jomanix (2022, *N* = 23; 2023, *N* = 2; 2024, *N* = 5). We recorded 17 hand-clap events during the behavioural observations and recorded 13 hand-clap events on video (see https://osf.io/4xqp9/?view_only=2bc8b44888644014acfff2ca125b687c).

For each hand-clap event, we recorded whether the gesture appeared to be directed at a target (i.e., a recipient of the gesture). If so, the identity of this partner, and any potential response of the latter to the hand-claps. We also recorded whether the hand-claps occurred in combination with any other behaviours or facial expressions from Jomanix. A behaviour or facial expression would be considered to occur in combination with a hand-clap if it preceded or followed the hand-clap within a same interaction episode. These data were extracted from the behavioural observations as well as coded from the video recordings.

In the case of the Tonkean macaque Wallace, data on the hand-clap events were not actively collected, but rather the reports rely on opportunistic observations of the gesture being displayed by the individual. Two events were recorded on camera, one in 2018 and the other in 2023, each year corresponding to one of two study periods at the Primate Centre of Strasbourg University.

## 3. RESULTS

In 29 of the 30 hand-clap events, a partner was recorded, i.e., the gesture was directed at the recipient. In three of these events, due to uncertainty regarding the specific recipient of the gesture, two partners were recorded, which resulted in 32 identified recipients of the gesture. The hand-claps were directed in 15 cases at a juvenile or subadult (1-6 years old; nine times at an unknown juvenile, and six times at the same subadult male), in 14 cases at an adult (7-19 years old; 13 times at four different females, and one time at a male), and in three cases at an old individual (>20 years old; three different males).

Out of the 30 recorded hand-clap events, 25 hand-claps occurred in combination with other behaviours or facial expressions from Jomanix. All of these were agonistic signals. More specifically, twelve hand-claps occurred in combination with an aggressive posture or gesture (e.g., lunge at partner, ground slap), six hand-claps occurred in combination with a threatening facial expression (e.g., stare, open mouth threat), and seven hand-claps occurred in combination with several agonistic signals displayed in a same interaction episode. Examples of Jomanix using the hand-clap in combination with other signals include him lunging at a partner and then hand-clapping (e.g., Video 1), him slightly moving his head and shoulders toward a partner, staring at the latter and then hand-clapping (e.g., Video 2), or various sequences of him displaying the stare or open mouth threat, the lunge and/or ground slap, along with the hand-clap (e.g., Videos 3, 4, and 5). In all instances, Jomanix was oriented towards and looking at the target of his hand-clap gesture.

For seven of the hand-clap events, no response from the recipient of the gesture could be recorded. This happened, for instance, when the recipient did not appear to see the hand-clap (e.g., Videos 6 and 7) or when the recipient was out of video shot and no potential response could be seen on film (e.g., Video 8). For all other hand-clap events, a response from the recipient was recorded. As in three events, two partners were recorded as potential recipients of the gesture, there was a total of 25 responses recorded across all hand-claps emitted. Recipients of the gesture responded in 16 cases with submissive behaviour or vocalisation (e.g., give ground, flee, squeaks) and in nine cases with counter-aggressive behaviour or vocalisation (e.g., lunge, scream face, head bob) (Fischer & Hammerschmidt, 2002; Hesler & Fischer, 2007).

After our second field season (2023), park staff members reported that another subject from the same group as Jomanix had been observed hand-clapping. The individual was an adult male, 12 years old at that time (i.e., born in 2011). He passed away before we returned to the park in 2024, and no hand-clap events could ever be recorded on video from him.

In the case of the Tonkean macaque, Wallace, the individual was only ever observed hand-clapping at humans. From the two hand-clap events caught on camera, the first one shows Wallace hand-clapping when an experimenter is on the other side of the enclosure mesh and in possession of food in her belt bag (Video 9), and the second one while he is participating in an experiment where the experimenters handle foods he can receive as rewards (Video 10). Thus, in both instances, the gesture was directed at human recipients, and occurs in the presence of food.

## 4. DISCUSSION

Our observations suggest that the hand-clap has a communicative function and constitutes an agonistic signal for Jomanix. He directs the gesture at specific recipients, mainly combines it with other agonistic behaviours and/or facial expressions, and generally elicits either a submissive or counter-aggressive response from the recipients.

Although we do not have direct evidence, it does not seem entirely unlikely that Jomanix acquired the gesture from the park staff. To shoo the monkeys away from the visitors, the park staff members coax the animals off the walking paths by vocalising, whistling, or walking towards the animals, and occasionally hand-clapping. The Barbary macaque Jomanix may thus have copied the hand-clapping from them, and using it in a similar way to signal to the recipient to move out of the way. The opportunity for captive animals to observe, and sometimes interact with, humans may promote occasional acquisition of novel gestures through spontaneous imitation (e.g., chimpanzees imitating zoo visitors in knocking on enclosure windows with knuckles or palm, Persson et al., 2018). Most often, however, inter-species imitation requires human actions to be actively demonstrated or taught to the animals (e.g., Byrne & Tanner, 2006; Call, 2001; Custance et al., 1995; Kumashiro et al., 2003). In addition, it is argued that even in cases of animals mimicking human actions, it is unclear whether they possess adequate understanding of the function and intention of the gestures to put the latter to use in achieving communicative goals (discussed in Byrne & Tanner, 2006). In the case of Jomanix, however, while the notion of him imitating staff members’ hand-clapping remains speculative, our observations suggest that he uses the gesture as a means to deter other monkeys, similarly to how the action is intended by the humans he observed.

While hand-clapping has been reported in other studies and across different species, the apparent use and function of the gesture is not always the same. Examples include chimpanzees’ hand-clapping with conspecifics during play (Liebal et al., 2004; Tomasello et al., 1994) or greetings (Scott, 2013). Bonobos hand-clap to elicit grooming (Ingmanson, 1987; de Waal, 1988), and gorillas hand-clap when vigilant and seemingly intending to alert conspecifics to danger (Fay, 1989; Salmi & Muñoz, 2020) or maintain group cohesiveness (Kalan & Rainey, 2009). In the case of Wallace, the Tonkean macaque described in this study, the circumstances in which he uses the hand-clap gesture contrast with those of Jomanix as well. Indeed, Wallace hand-claps seemingly to attract the attention of human keepers and experimenters in the presence of food. Captive primates have been observed learning gestures directed at humans such as hand-clapping for attention (Fletcher, 2006; Hostetter et al., 2001) or to solicit assistance during out-of-reach foraging tasks (Cartmill & Byrne, 2007), or pointing to request food (Leavens & Hopkins, 1999). These examples may underline the role of captivity, i.e., human endorsement and/or potential constraints of the setting, in the occasional development of novel gestures to fulfil a certain goal (discussed in Hobaiter & Byrne, 2011). Similarly, our reported case of hand-clapping in another macaque, displayed in an entirely different context, supports the notion that each of the gestures are associated with a specific function (displace vs attention-getting), and that this function may be socially shaped in accordance with the surrounding human influence.

To date, only one other monkey from Jomanix’s group was reported replicating the gesture since Jomanix was first seen to display it, and in that case perhaps through human facilitation also. Alternatively, it is suggested that gestural acquisition may possibly occur via observation of the gesture in question being used by another conspecific (Liebal & Call, 2012; Pika & Fröhlich, 2019), but it is unclear whether and to what extent this phenomenon occurs in primates. Theory suggests that this process would result in a high degree of repertoire concordance within a group (Call & Tomasello, 2007; Prieur et al., 2020), yet the hand-clap gesture has not, with the exception of one other individual, spread within Jomanix’s group thus far. Taken together, our results corroborate the notion that gestural acquisition by social transmission may occur in monkeys as well as in great apes (e.g., chimpanzees’ grooming hand clasp, Bonnie & de Waal, 2006; McGrew & Tutin, 1978) but the frequency with which it happens is extremely low (Amici & Liebal, 2023; Dean et al., 2018; Prieur et al., 2020).

While it is unclear whether it is the hand-clap itself that the recipients react to, Jomanix is able to flexibly combine the gesture with other established agonistic signals to achieve a tangible response from his targets. Jomanix’s hand-clap gesture can be classified as a case of multimodal signalling where signals encompass different sensory modalities (Rowe, 1999), by combining a visual (e.g., head bob, lunge) with an auditory (hand-clap) gesture. In addition, the behaviour can be conceived as multi-component signalling, where different signal components are integrated regardless of their modality (Micheletta et al., 2013), such as combining a facial expression (e.g., stare, open mouth threat) with a gesture (hand-clap). Suggested functions of signal combinations include amplification, modulation and refinement of signal meaning, and ultimately could permit greater communicative flexibility and complexity (Fröhlich & van Schaik, 2018; Pollick & de Waal, 2007). In addition, the use of the hand-clap for Jomanix is clearly goal-directed and fulfils the criteria for first-order intentional communication, which requires that the signaller intends to signal to produce a change in the behaviour of the recipient (Fischer & Price, 2017; Townsend et al., 2017). Jomanix therefore demonstrates flexibility and first-order intentionality, two hallmarks of gestural communication in great apes (Byrne et al., 2017; Call, 2008).

There is discussion relative to the possibility that idiosyncratic gestures may be artefacts of insufficient sampling and actually constitute behaviours of the larger species’ repertoire that (simply) haven’t been observed yet (Byrne et al., 2017; Genty et al., 2009; Hobaiter & Byrne, 2011), but this notion seems unlikely in our case. The population of Barbary macaques at ‘La Forêt des Singes’ has been monitored since the park’s establishment in 1974 (de Turckheim & Merz, 1984), and actively studied for various research projects since the 1990’s (e.g., Almeling et al., 2017; Bosshard et al., 2024; Fischer, 1998; Fischer & Hammerschmidt, 2001; Hammerschmidt et al., 1994; Patzelt et al., 2009; Rathke et al., 2022), yet hand-clapping was never reported before – nor has the gesture been reported in other populations of the same species. While we cannot rule out a certain predisposition of the species to develop a given gesture, our observations rather imply some form of social facilitation from humans in the appearance or acquisition of novel gestures. This idea is supported by the fact that we report two different instances of hand-clapping from macaques in two different populations, where the form of the gesture may have been learned from humans, but the function developed according to each individual’s particular intention or need.

This case study adds to the body of evidence demonstrating that non-ape primates are able to produce, develop and use novel gestures with first-order intention and flexibility. Thus, novel gesture acquisition may be a more common phenomenon in monkeys than previously assumed. In addition, reports such as the present one may mark the inception of a gesture within a group or population, help in reconstructing the diffusion process of the gesture, and broaden our knowledge relative to the spectrum of communicative and contextual use of the gesture within a species.

## ACKNOWLEDGEMENTS

We thank Ellen Merz, Gilbert de Turckheim, and Roland Hilgartner for their permission to conduct our observations at ‘La Forêt des Singes’. We are also particularly grateful to the staff members of the park, who were dedicated to helping us catch the hand-clap events on camera, provided us with additional information relative to Jomanix’s hand-clapping history, and sent us one of the hand-clap videos. We also thank Victor Gass, Nadja Vögtle, and Claire des Pallières for their assistance in data collection in the field, and Dr. Derek Murphy for his aid in filming some of the hand-clap events. Finally, we thank the University of Strasbourg and Silabe (https://www.silabe.com/) for supporting data collection on the Tonkean macaques and providing animal care.

## AUTHOR CONTRIBUTIONS

**Tiffany C. Bosshard:** Conceptualization; Data curation; Formal analysis; Investigation; Project administration; Visualization; Writing – original draft preparation; Writing – review & editing. **Marie Hirel:** Writing – review & editing. **Hélène Meunier:** Resources; Writing – review & editing. **Julia Fischer:** Conceptualization; Funding acquisition; Project administration; Resources; Supervision; Writing – review & editing.

## DATA AVAILABILITY STATEMENT

The data associated with this article can be found online at https://osf.io/4xqp9/?view_only=2bc8b44888644014acfff2ca125b687c

